# Mechanosensing activates flashing Ca^2+^ dynamics associated with cell regeneration in *Physcomitrium patens*

**DOI:** 10.64898/2026.07.26.740844

**Authors:** Hong-An Wang, Yun-Ching Yen, Han Tang

## Abstract

Mechanical wounding is the primary trigger for various forms of plant regeneration; however, the mechanism by which mechanical cues are rapidly translated into regenerative cell fate decisions remains elusive. Here, using live-cell imaging in *Physcomitrium patens*, we established a spatiotemporal framework of Ca^2+^ signaling that bridges initial wound perception and cellular reprogramming. We demonstrate that wounding induces a primary Ca^2+^ wave followed by two forms of secondary responses, including a unique stochastic flashing signature that persists for hours and restricts in wound-neighbor cells. Wound-induced membrane deformation was associated with the activation of Ca^2+^ influx. In addition, osmotic stress-induced membrane tension also elicited a rapid Ca^2+^ spike followed by broadly flashing signals without spatial specificity. Pharmacological blockade of plasma membrane Ca^2+^ channels and chelation of extracellular Ca^2+^ significantly reduced both primary and secondary Ca^2+^ responses, indicating that these wound-induced Ca^2+^ dynamics mainly rely on regulated influx rather than passive diffusion. Inhibition of Ca^2+^ signaling also disrupted mechanosensitive F-actin reorganization and suppressed cell reprogramming, resulting in a severe reduction in regeneration capacity. Together, our results demonstrate a complex spatiotemporal Ca^2+^ code that precedes cytoskeletal reorganization and cell reprogramming in wound-neighbor cells, providing new insights into plant tissue regeneration.

**Highlight:** Mechanical stress triggers a rapid Ca^2+^ influx followed by a prolonged flash signaling, revealing a spatiotemporal framework of Ca^2+^ dynamics that links membrane tension to coordinated plant regeneration.

## Introduction

Regeneration is an essential process that restores wounded tissues or replaces lost organs, a phenomenon found in both animals and plants (Birnbaum and Alvarado, 2008). Among distinct forms of plant regeneration, mechanical wounding is a universal primary trigger. Upon wounding, the adjacent cells sense the mechanical stimuli, consisting of biochemical cues and mechanical cues, that spread from damaged loci and are converted into subcellular signal transduction cascades that ultimately trigger cell reprogramming and regeneration (Ikeuchi *et al*., 2019; Sugimoto *et al*., 2019). The perception of mechanical cues and subsequent cellular responses in plants involves a rapid activation of wound signals such as calcium (Ca^2+^) spikes (Koselski *et al*., 2023; Tian *et al*., 2020).

Such a rapid and transient increase in cytosolic Ca^2+^ concentration ([Ca^2+^]cyt) is an early event in many mechanosensory responses, including touch, osmotic stress, and gravitropism (Suda and Toyota, 2022). These Ca^2+^ responses are not uniform; they form specific “calcium signatures” characterized by variations in amplitude, duration, frequency, and localization that encode information about the nature of the mechanical stimulus (Suda and Toyota, 2022). Given that apoplastic Ca^2+^ concentrations are approximately three orders of magnitude higher than cytosolic [Ca^2+^], mechanosensitive (MS) ion channels at the plasma membrane are thought to contribute directly to Ca^2+^ influx (Guichard *et al*., 2022). Further downstream, Ca^2+^ also plays a role in activating genes associated with stress responses (Li *et al*., 2019b; Tian *et al*., 2020). The regulation of intracellular Ca^2+^ levels involves influx from the apoplastic space and internal stores, e.g., the vacuole and endoplasmic reticulum, and efflux systems that remove Ca^2+^ from the cytoplasm (Tian *et al*., 2020).

Mechanical wounding perturbs the cell wall-plasma membrane-cytoskeleton continuum, which provides a physical linkage between the extracellular environment and intracellular signaling (Bacete and Hamann, 2020). Upon wounding, microtubules have been shown to reorganize and align with the direction of cellular tension in the shoot apical meristem of *Arabidopsis* (Hamant et al., 2008). In Arabidopsis root, laser ablation induced cell damage results in microtubules stabilization in wound-neighbor cell that ultimately regulate the orientation of regenerative cell division plane (Hoermayer *et al*., 2024). Despite numerous studies found cytoskeletal reorganization in response to wounding, F-actin responses are less clearly defined and have not been observed as consistently as those of microtubules.

In our previous research, we demonstrated that in an excised leaf from a gametophore of *Physcomitrium patens (P. patens)*, F-actin rapidly accumulates at the cortical membrane of cells adjacent to the wounding site, while no such accumulation appears in the opposite membrane (Huang *et al*., 2025). In tip-growing cells, elevated Ca^2+^ can inhibit apical actin accumulation, while low Ca^2+^ promotes F-actin assembly (Cárdenas *et al*., 2008). Disrupting actin can also affect Ca^2+^ oscillation profiles (Bascom Jr *et al*., 2018). Whether the Ca^2+^ signaling plays a role in wound-induced F-actin reorganization remains unclear.

Upon wounding or touch, although numerous studies have shown a rapid Ca^2+^ wave in different species as an early wound response (Storti *et al*., 2018b; Watanabe *et al*., 2024; Zhou *et al*., 2025), how Ca²⁺ signals evolve beyond the initial response, and how individual cells interpret these signals, remain less well understood. Moreover, the relationship between distinct Ca^2+^ dynamic patterns and downstream outcomes, such as cellular reprogramming or regeneration, remains largely unresolved. A spatiotemporal framework for interpreting wound-induced Ca^2+^ signaling beyond its initial activation is lacking.

To resolve the spatiotemporal framework of wound-induced Ca^2+^ signaling at single-cell resolution and assess its relationship with cytoskeletal remodeling and regeneration, we employed the moss *P. patens* as a model system.

*P. patens* is characterized by a high capacity for regeneration. When a moss leaf is excised from a gametophore, leaf cells adjacent to the wound can reprogram into stem-cell-like, filamentous tissue called protonemata and regenerate a new filament from a single leaf cell. Alternatively, some wound-adjacent leaf cells can re-enter the cell cycle to undergo regenerative division (Ishikawa *et al*., 2011). A few master regulators, e.g., STEMIN and WOX13, have been identified as initiators of reprogramming through chromatin remodeling (Ishikawa *et al*., 2019; Sakakibara *et al*., 2014), while how they are activated by wounding stimuli remains unclear.

In this study, we utilized precise laser ablation in *P. patens* to investigate the spatiotemporal dynamics of wound-induced Ca^2+^ signaling at single-cell resolution. We identified a rapid primary Ca^2+^ wave followed by distinct secondary responses, most notably a stochastic flashing Ca^2+^ signature that persists for hours, specifically in wound-neighbor cells destined for reprogramming. Plasma membrane deformation induced by physical damage or osmotic stress can sufficiently elicit a rapid Ca^2+^ spike and prolonged flashing signals. Further, the wound-induced prolonged flashing Ca^2+^ signaling is essential for downstream F-actin reorganization and successful cell fate transition. By integrating physical mechanosensing with long-term physiological responses, our work reveals a unique Ca^2+^-mediated mechanism that translates immediate wound impacts into regenerative outcomes.

## Results

### Wounding induces a rapid and biphasic Ca²⁺ signaling response

To investigate the wound-induced Ca^2+^ signaling response in *P. patens* and determine whether it exhibits conserved patterns observed in other plant species, we performed high-resolution live imaging following laser ablation and monitored cytosolic Ca^2+^ dynamics using the genetically encoded Ca^2+^ indicator GCaMP3 (Kleist *et al*., 2017). Live-cell imaging revealed that wounding elicited a rapid and biphasic Ca^2+^ response. Immediately following ablation, a transient primary Ca^2+^ spike was induced at wound-neighbor cells and then propagated radially into wound non-neighboring cells and gradually dispersed over the time course of approximately 10-15 minutes (**Fig. 1A**). The Ca^2+^ wave diminished progressively, disappearing first in wound-neighbor cells and subsequently in more distal cells, mirroring the sequence of signal propagation (N >200 ablation sites, wave occurrence rate: 100%. **Fig. 1A, Video S1**).

**Figure 1.**
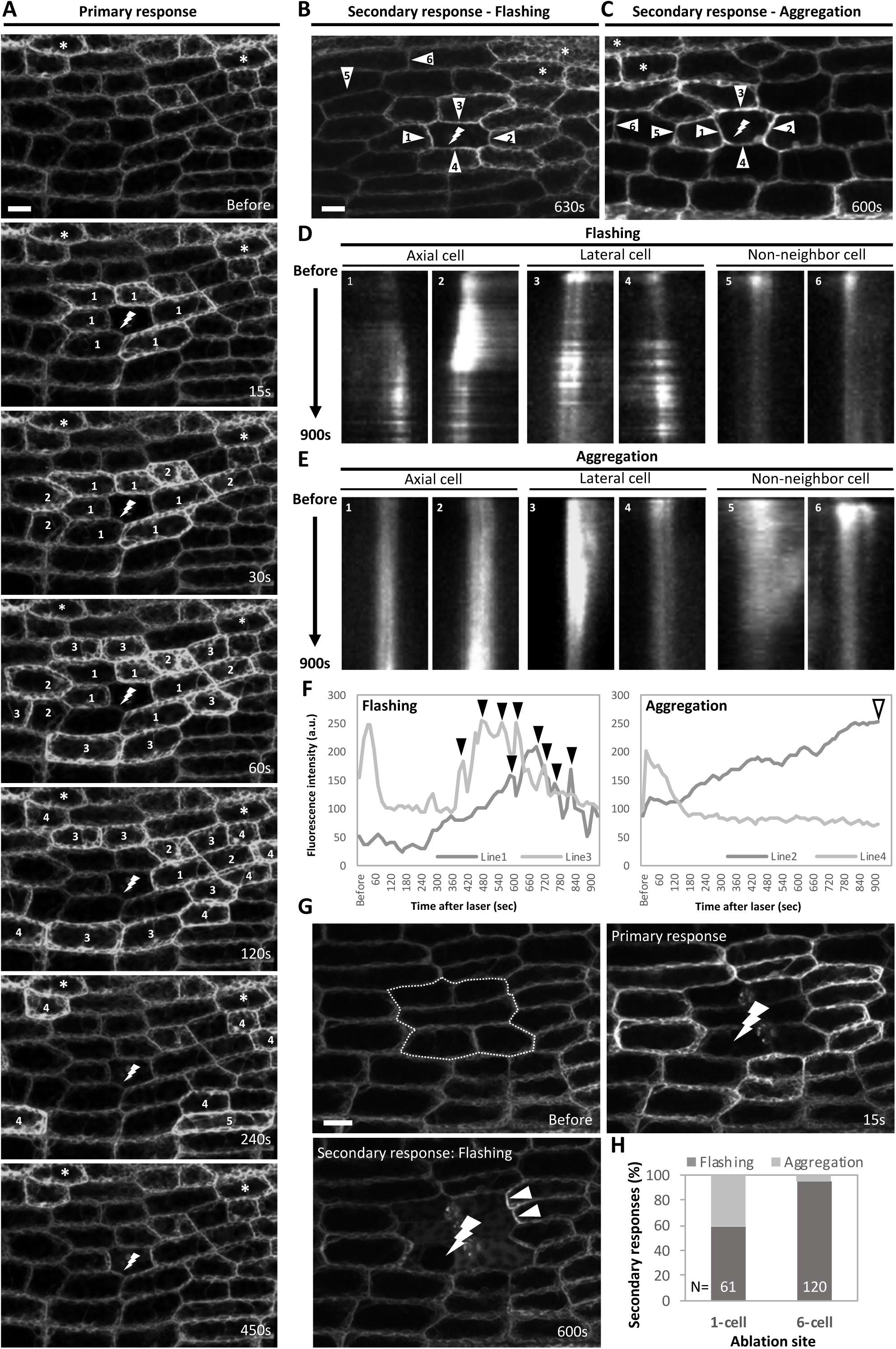
Wounding induces a rapid and biphasic Ca^2+^ signaling response. **(A)** Representative images showing the primary Ca^2+^ response induced by single-cell laser ablation in *P. patens* leaves expressing GCaMP3. Images were acquired at the indicated time points before and after ablation. Laser-targeted cells are marked by lightning symbols. Numbers denote cells exhibiting Ca^2+^ activation at successive time points following wounding. ‘1’ indicates cells activated at 15 seconds (s), ‘2’ at 30 s, and ‘3–5’ indicates cells activated between 60 and 240 s. Number labels were removed once Ca^2+^ signals returned to baseline. Asterisks mark cells displaying Ca^2+^ signals that did not respond to wounding. At 450 s, the primary Ca^2+^ signals diminished to baseline, as shown before wounding. Scale bar = 20 µm. **(B)** and **(C)** Representative images at later time points showing secondary Ca^2+^ flashing and aggregation activities in wound-neighbor cells. Numbered arrowheads point to the membranes used for kymograph analysis in **(D)** and **(E)**. Scale bar = 20 µm. **(D)** and **(E)** Kymographs of Ca^2+^ fluorescence along membrane-crossing (mc) lines corresponding to **(B)** and **(C)**. Axial and lateral wound-neighbor cells are shown separately, with non-neighbor cells included as internal controls. **(F)** Quantification of Ca^2+^ intensity from the kymographs in **(D)** and **(E)**. For flashing, the axial cell (kymograph line 1) and the lateral cell (kymograph line 3) obtained from **(D)** are compared in the left panel. Multiple Ca^2+^ peaks that show up at random timepoints with varying intensity (arrowheads) are defined as flashing activity. For aggregation, the axial cell (kymograph line 2) and the lateral cell (kymograph line 4) obtained from **(E)** are compared in the right panel. Progressive Ca^2+^ accumulation specifically at the wound-adjacent membrane of axial cells, pointed by a hollow arrowhead, is defined as aggregation activity. **(G)** Representative images showing the primary Ca^2+^ response and secondary flashing activity induced by a six-cell laser ablation. The laser target area is depicted by a dotted frame before the laser and a lightning symbol after the laser. Arrowheads denote cells with flashing signals. Scale bar = 20 µm. **(H)** The proportion of events that occurred with flashing or aggregation patterns under 1-cell and 6-cell ablation.

Following the primary response, two types of secondary Ca^2+^ activity were observed in wound-neighbor cells: flashing and aggregation (**Fig. 1B–1F, Video S2, S3**). To examine the dynamics of the Ca^2+^ signals, we measured the intensity of the Ca^2+^ indicator over time at wound-neighbor cells and non-neighbor cells. Intensity along a line drawn across the indicated membrane with number marks in **Fig. 1B, 1C** was measured for kymograph analysis, as shown in **Fig. 1D, 1E**. A segmented line along the signals shown in the kymograph was measured over time as shown in **Fig. 1F**. In the flashing response, wound-neighbor cells exhibited random Ca^2+^ signaling spikes that frequently, though not necessarily, localized near the plasma membrane adjacent to the wound. These flashes occurred with irregular timing and frequency on each detected side of wound-neighbor cells (N > 200 ablation sites, **Fig. 1D, 1F**). In contrast, non-neighboring cells showed only the primary Ca^2+^ response and did not exhibit secondary activity (**Fig. 1D, Video S2**). Notably, due to the rapid response of the primary wave, the wave signal might be missed in the kymograph analysis of wound-neighbor cells, as shown in **Fig. 1D, 1E**, e.g., line 1, in **1D** and line 1 in **1E**. We reasoned that the primary wave signal passed through the measured area within the first 15 seconds.

Aggregation is the other type of secondary response, characterized by sustained Ca^2+^ accumulation at the plasma membrane of wound-neighbor cells aligned along the wound axis, whereas lateral neighboring cells did not show comparable signal enrichment (N> 200 ablation sites. **Fig. 1E, 1D**). The signal accumulated at lateral cells diminished within 900 seconds, e.g., line 3 in **1E**, while the aggregation of Ca^2+^ signals at axial cells remained over the examined timeframe.

We identified distinct Ca^2+^ signaling dynamics in response to wounding. We next tested whether these patterns are associated with differences in wound size or severity. To achieve this, we observed and compared Ca^2+^ signaling following single- or six-cell ablation (**Fig. 1G, Video S4**). We counted the number of ablation areas exhibiting flashing or aggregation of Ca^2+^ signals by visual confirmation of each video. We found that when a single cell was ablated, flashing and aggregation responses occurred at approximately equal frequencies (N=61 ablation sites, **Fig. 1H**). However, increasing the wound size to a six-cell area led to a strong bias toward flashing responses, observed in approximately 95% of wounded regions (N=120 ablation sites, **Fig. 1H**). This difference indicates that the spatial patterns and dynamics of secondary Ca^2+^ signaling are dependent on wound size and severity.

### The Ca^2+^ flashing signal persist for hours in wound-neighbor cells

To examine the duration of the secondary Ca^2+^ flashing signal, we performed long-term time-course imaging. Remarkably, the presence of Ca^2+^ flashing signals remained detectable for up to 8-16 hours after wounding, although the magnitude and frequency of flashing gradually declined over time (**Fig. 2**). In addition to the frequency, the number of wound-neighbor cells that exhibited Ca^2+^ flashing signals also decreased over time (Fig. **2I**). To our knowledge, wound-induced Ca^2+^ signaling appears to be transient, lasting only for a few minutes, while such prolonged wound-induced Ca^2+^ signaling has not been reported previously.

**Figure 2.**
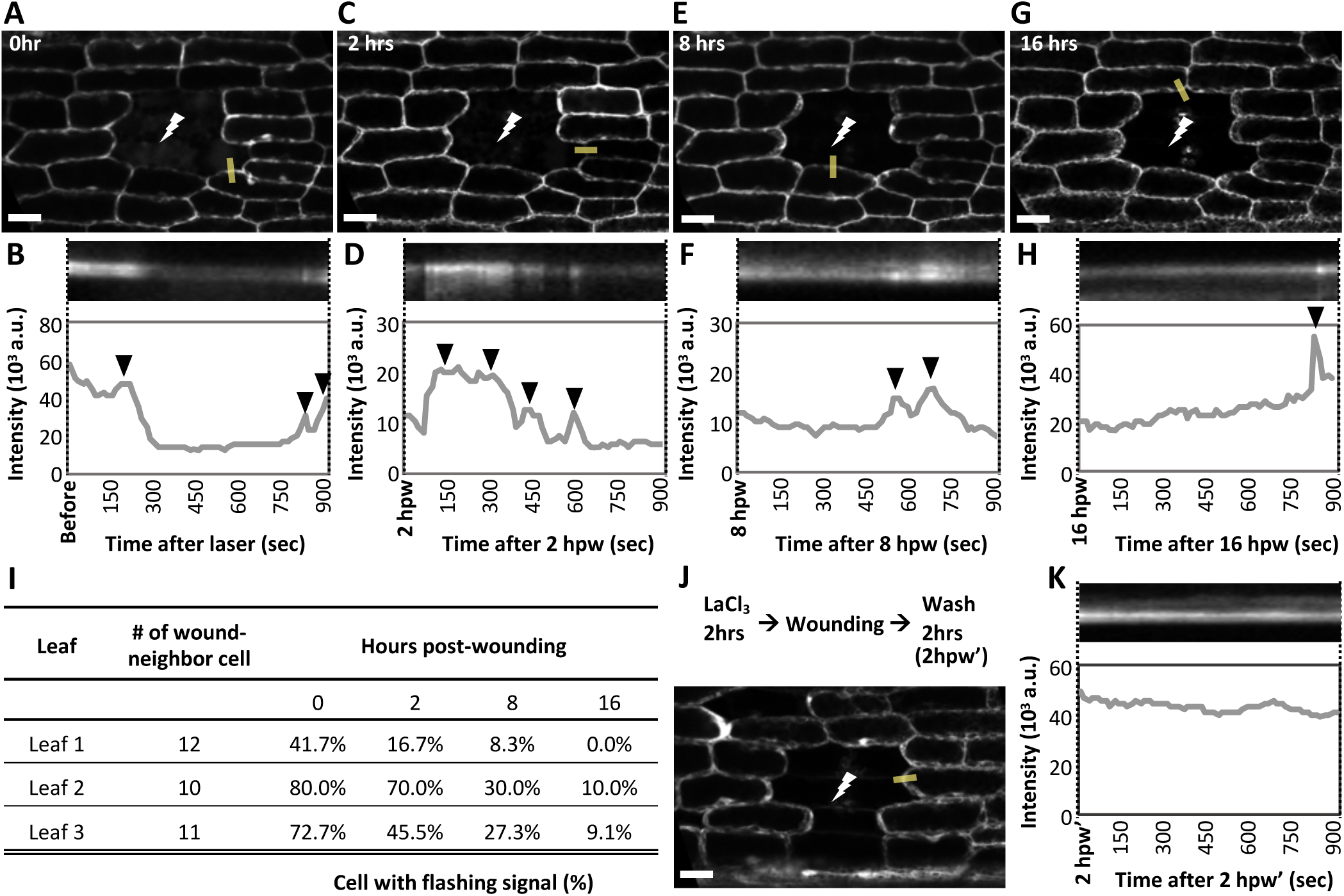
The Ca^2+^ flashing signal persists for hours in wound-neighbor cells. Excised leaves from the GCaMP3 line were ablated and placed on agar medium for a long-term observation. The leaf was moved to a slide for a time-lapse imaging at 0, 2, 8, and 16 hours post-wounding (hpw). The time-lapse imaging was acquired under a confocal microscope with a 15-second interval and continued for 900 seconds. **(A) (C) (E) (G)** The representative images of the same wound site at 0, 2, 8, and 16 hpw. Examples of flashing Ca^2+^ signals at each time point were visually selected and depicted by yellow lines drawn across different membrane regions. Scale bar = 20 µm. **(B) (D) (F) (H)** Kymograph analyses along the yellow lines are shown; dotted lines align the kymographs with the corresponding time points. The dynamics of flashing signals were measured by a line-drawing tool and presented in plot profiles, with arrowheads pointing to the random flashing signals. **(I)** Proportion of wound-neighbor cells that exhibit sustained flashing signals on 0, 2, 8, and 16 hours post-wounding. Three independent experiments. **(J)** Detection of Ca^2+^ signal at 2 hours post-washing (2hpw’) from LaCl_3_ treatment. The excised leaf was pre-treated with 5mM LaCl_3_ for 2 hours, followed by laser ablation under LaCl_3_ conditions, and washed in ddH_2_O for another 2 hours. **(K)** Kymograph analysis as described above. The Ca^2+^ signal was examined at 2hpw’ for 900s. The flashing Ca^2+^ signal was not observed. N=5 independent experiments; cell number > 25.

In addition to the restricted area of the secondary Ca^2+^ signaling at wound-neighboring cells, we verified that the flashing patterns of the Ca^2+^ signal are indeed triggered by wounding. We examined the cytosolic Ca^2+^ in unwounded leaves and focused on the intact leaf area and the cutting edge. Consistent with the laser ablation, the cutting edge appeared to show flashing signals (**Video S5**). Besides the cutting edge, the Ca^2+^ signal displayed a steady state as the baseline of cytoplasmic Ca^2+^, and no wave or flashing patterns were observed in 3 independent excised leaves (**Video S6**).

To test whether the secondary Ca^2+^ responses depend on the primary wave, we transiently blocked Ca^2+^ influx using Lanthanum Chloride (LaCl_3_), which has been widely employed to inhibit plasma membrane Ca^2+^ channels in plants (Li et al., 2019a). Excised leaves were pretreated with LaCl₃ for 2 hours and wounded in the presence of LaCl₃. Under these conditions, both the primary Ca^2+^ wave and subsequent flashing activity were significantly suppressed, as showed in the following results (**Fig. 5**). The treated leaves were then washed and allowed to recover from inhibition for 2 hours, followed by a second time-lapse imaging session at the same regions to monitor secondary flashing signals. We reasoned that, if activation of the secondary signal depends on the primary signal, inhibition of the primary response should suppress secondary flashing even after washout. Indeed, no obvious secondary flashing signals were observed, whereas the overall Ca^2+^ intensity in wound-neighbor cells increased, suggesting a disruption of intracellular Ca^2+^ homeostasis (N=4 independent experiments; cell number>25; **Fig. 2J, 2K, S6**). Taken together, these results define a new biphasic Ca^2+^ signaling response to wounding, consisting of a rapid, transient primary wave followed by prolonged secondary responses with distinct signatures.

### The wound-induced plasma membrane deformation contributes to the activation of Ca^2+^ signaling

Previous studies have shown that Ca^2+^ signaling can be activated by mechanical forces that deform the plasma membrane, thereby opening mechanosensitive channels and allowing Ca^2+^ influx (Fan *et al*., 2004; He *et al*., 2018; Kurusu *et al*., 2013). To test whether wound-induced Ca^2+^ signaling is activated by mechanical changes rather than cellular damage *per se*, we examined plasma membrane deformation and assayed Ca^2+^ signal induction. We applied laser ablation at different power levels to uncouple membrane deformation from cell damage. Ablation at full laser power (100%) induced a robust Ca^2+^ response and caused outward deformation of the wound-adjacent plasma membrane, bowing into the ablated cell (**Fig. 3A**), while the ablated cell was killed and shrank. The cell death was confirmed by cell content leakage, induction of autofluorescence, and cell shrinkage, which has been verified in our previous study (Huang *et al*., 2025). 450 seconds after ablation, the cell membrane deformation was demonstrated using cell outline staining and overlapping images from before and after ablation (**Fig. 3A**). With full laser power, the membrane adjacent to the ablated cell before ablation did not overlap with the membrane after ablation, indicating a membrane displacement. Under this condition, with wound-induced membrane deformation, we consistently observed Ca^2+^ responses as described before (N> 60 independent experiments).

**Figure 3.**
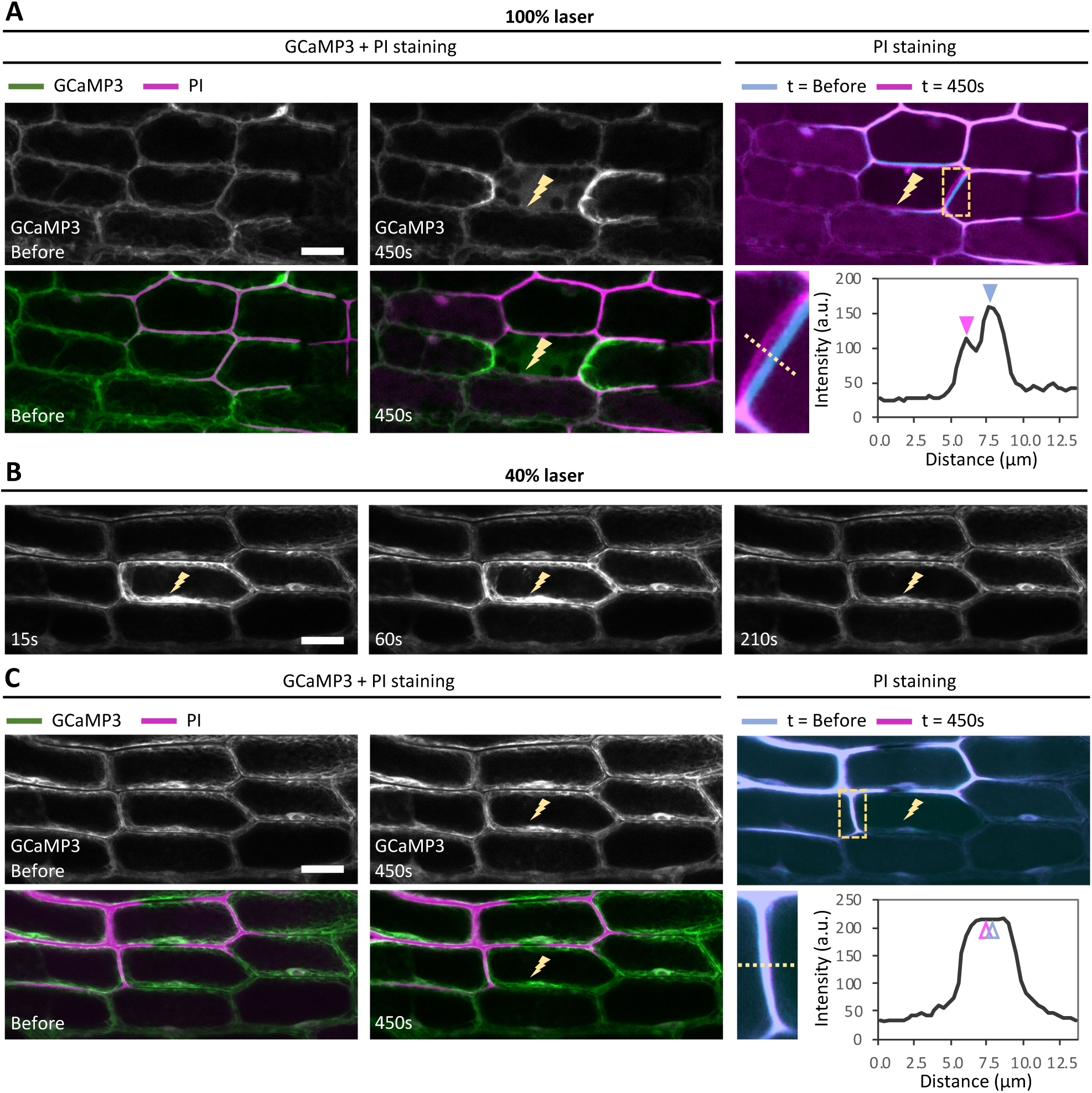
Plasma membrane deformation is required for the wound-induced Ca^2+^ signaling. **(A)** Representative images show the aggregation of the Ca^2+^ signal tracing the cell outline stained with propidium iodide (PI staining). With a full-power laser, the Ca^2+^ signal is aggregated in axial cells adjacent to the ablated cell (lightning symbol). The cell outline images before and after ablation are overlaid in the right panel. The signal for t=before was color-coded with cyan; the signal for t=450 seconds was color-coded with magenta. The hashed frame depicts the interface between the neighbor and the ablated cell, enlarged in the panel below. The intensity along the dotted line was measured and showed two peaks (magenta and cyan arrowheads), demonstrating plasma membrane deformation induced by laser ablation. N= 61 independent experiments. **(B)** and **(C)** Example images present the transient response of Ca^2+^ signals in the laser-targeted cell with a low-power laser (40%), N=3 independent experiments. **(B)** The ablated cell exhibits a transient Ca^2+^ signal, while no primary or secondary responses were observed in neighboring cells. **(C)** The cell outline images before and after ablation are overlaid in the right panel. The hashed frame and dotted line were presented in the same way as described in **(A)**. The hollow arrowheads point to the overlap of cyan and magenta signals, demonstrating that no membrane deformation is caused by 40% laser ablation. Scale bar = 20 µm.

In contrast, ablation at lower power (40%) damaged the target cell that exhibited a transient Ca^2+^ spike, which diminished within 210 seconds (**Fig. 3B**), while no displacement in cell shape or plasma membrane was observed (**Fig. 3C, right**). In wound-neighbor cells, the Ca^2+^ baseline remained the same, but no Ca^2+^ primary wave or secondary activities were detected (**Fig. 3C, left,** N=3 independent experiments). In sum, these results suggest that plasma membrane deformation is associated with the activation of wound-induced Ca^2+^ signaling, supporting a mechanosensitive origin of this response.

### Osmotic stress-induced membrane tension triggers a rapid Ca^2+^ signaling

We next asked whether plasma membrane deformation or tension alone is sufficient to induce Ca^2+^ signaling in the absence of cell damage. To test this, we applied mannitol to impose hyper- or hypo-osmotic stress, thereby altering cell volume and membrane tension without physical wounding.

Given the rapid kinetics of Ca^2+^ signaling, we used a microfluidic device that enables unidirectional liquid flow through sample leaves and manual exchange of the surrounding medium (**Fig. 4A**). Live-cell imaging revealed that mannitol application triggered a rapid Ca^2+^ wave that propagated in the same direction as the fluid flow. This response was obvious but transient, initial observations after 50 seconds but dispersing within approximately 2 minutes (**Fig. 4B**). After prolonged mannitol treatment (∼20 minutes), clear plasmolysis was observed, indicating sustained hyper-osmotic stress (**Fig. 4C**). Under these conditions, subsequent laser ablation of a single cell failed to induce visible plasma membrane deformation in neighboring cells (**Fig. S1**).

**Figure 4.**
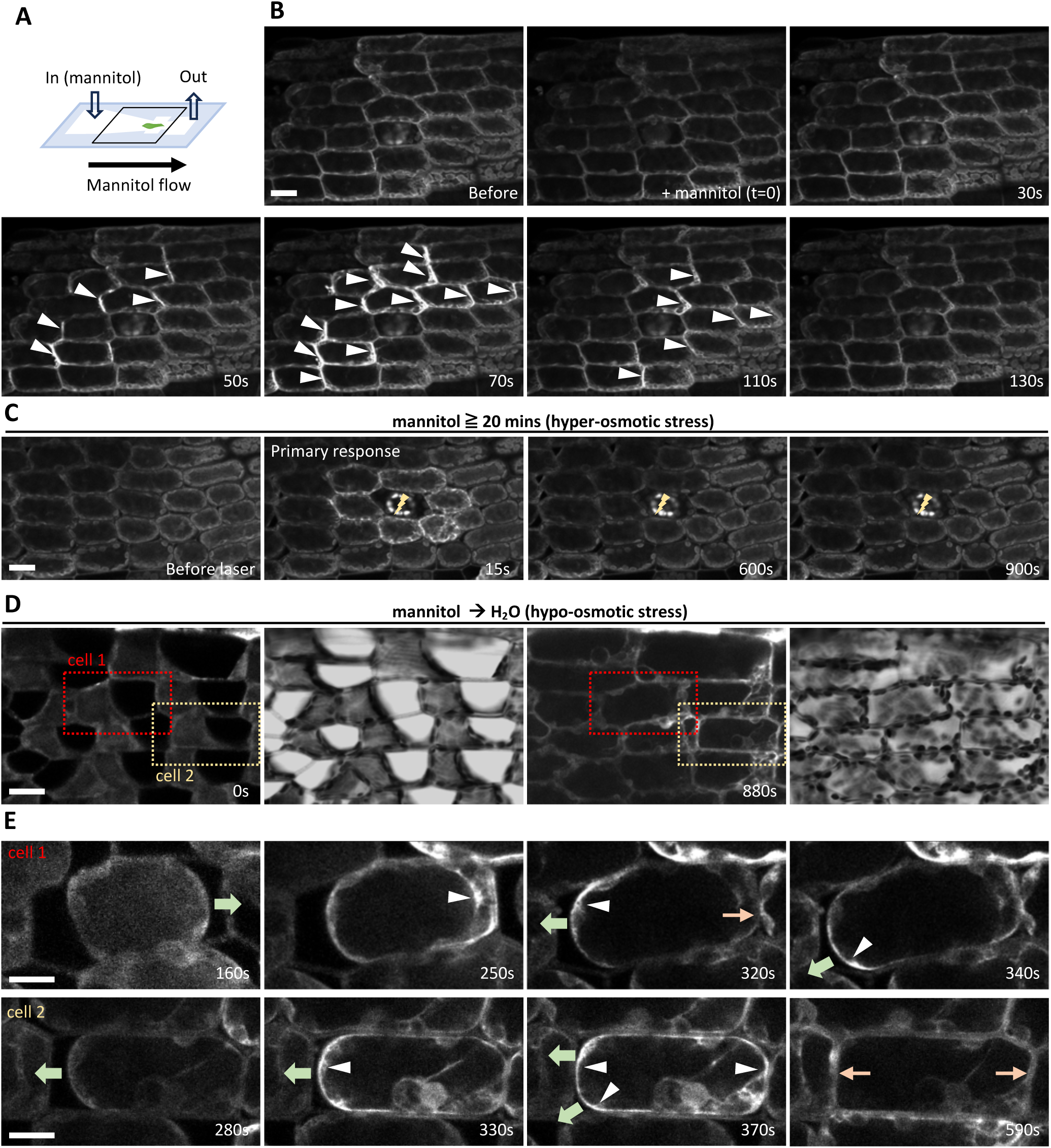
Membrane tension is sufficient and necessary to activate wound-responsive Ca^2+^ signaling. **(A)** Schematic of the microfluidic setup used for osmotic manipulation. A single leaf from the GCaMP3-expressing line was placed in a home-made microfluidic chamber and covered with a coverslip, leaving an inlet and an outlet for rapid medium exchange (white arrows). Hyper-osmotic treatment was applied by flowing 700 mM mannitol through the inlet; flow direction is indicated by the black arrow. **(B)** Time-lapse images acquired at 15-second intervals following mannitol application (time 0). A rapid Ca^2+^ wave was induced upon hyper-osmotic treatment and propagated in the direction of mannitol flow, as indicated by white arrowheads. Scale bar = 20 µm. **(C)** Ca^2+^ signaling following laser ablation under hyper-osmotic conditions. Leaves were pre-treated with mannitol to induce plasmolysis, as confirmed by cell shape changes. Subsequent full-power laser ablation induced only a weak primary Ca^2+^ response and failed to trigger secondary Ca^2+^ activity. The ablated cell is marked by a lightning symbol. Scale bar = 20 µm. N=8 independent experiments. **(D)** Hypo-osmotic induction of plasma membrane tension. The surrounding medium was rapidly exchanged from 700 mM mannitol to water, inducing hypo-osmotic stress and outward plasma membrane extension. Under hyper-osmotic stress, no active Ca^2+^ signals are detected, and cells are shrunken to one side of a cell (0 sec). After 880 seconds, cells recover from hyper- to hypo-osmotic stress and subsequently refill the intracellular space. N=3 independent experiments. Two representative cells (outlined by dotted boxes and labeled cell 1 and cell 2) are shown in **(E)**. Scale bar = 20 µm. **(E)** Plasma membrane extension induces Ca^2+^ signaling. Transient Ca^2+^ signals were specifically enriched at regions of membrane extension and rapidly diminished once the plasma membrane (PM) reattached to the cell wall (CW). Green arrows demonstrate the direction of PM extension, white arrowheads point to the enriched Ca^2+^ signals, and orange arrows point to the attachment of PM to CW. Scale bar = 10 µm.

Consistent with suppressed membrane deformation, only a weak primary Ca^2+^ response was occasionally detected in response to wounding, and no secondary Ca^2+^ activity was observed, measured until 900 seconds after laser treatment (N=8 independent ablation, 3 out of 8 exhibited weak primary response, **Fig. 4C**). These results indicate that hyper-osmotic stress suppresses wound-induced Ca^2+^ signaling, likely by preventing plasma membrane deformation.

To test whether augmented membrane tension is sufficient to activate Ca^2+^ signaling, we replaced mannitol with water in the microfluidic device, thereby inducing a rapid transition from hyper- to hypo-osmotic conditions that mimics the outward membrane expansion observed in wound-adjacent cells (**Fig. 4D, 4E, Video S7**). During the recovery from hyper-osmotic stress, cell shapes and plasma membrane recovered from shrinkage to normal status (**Fig. 4D**). Along with the cell recovery, Ca^2+^ signaling was specifically enriched at regions of plasma membrane extension (**Fig. 4E, Video S7**). Notably, the Ca^2+^ signaling was not observed immediately when water flow approached the mannitol solution, but when shrunken cells extended their membrane and refilled the cellular space. This observation supported that Ca^2+^ activation is mediated through the membrane extension rather than basic liquid flow. Additionally, the spatial distribution of Ca^2+^ signals perfectly matched the direction of membrane expansion and gradually decreased once the plasma membrane reattached to the cell wall (N=3 independent experiments, **Fig. 4E, Video S7**). After the reattachment, the Ca^2+^ spike gradually returned to a baseline, followed by random flashing signals appeared across excised leaves lasting for 8 hours (N=5 independent experiments, **Video S8**).

These results demonstrate that plasma membrane tension is necessary and might be sufficient to activate Ca^2+^ signaling, supporting a mechanosensitive mechanism underlying wound-induced Ca^2+^ responses. While both localized membrane deformation induced by wounding and osmotic stress-induced membrane tension triggered prolonged Ca^2+^ responses, the resulting signaling patterns were different. Wounding induced spatially restricted Ca^2+^ flashing in wound-adjacent cells (**Fig. 2**), whereas osmotic stress elicited a more broadly distributed Ca^2+^ signaling (**Fig. 4, Video S8**). These observations suggest that cells might be able to distinguish between different modes of mechanical perturbation and generate stimulus-specific Ca^2+^ dynamic signatures. Such differences in spatiotemporal Ca^2+^ patterns may encode distinct downstream responses to localized injury versus global osmotic stress.

### Wound-induced Ca^2+^ signaling requires regulated Ca^2+^ influx across the plasma membrane

To further investigate the mechanism underlying wound-induced Ca^2+^ influx, we examined the effects of Ca^2+^ influx inhibitors on Ca^2+^ signaling following wounding. We applied two pharmacological treatments that interfere with extracellular Ca^2+^ entry through distinct mechanisms. First, we used the non-selective Ca^2+^ channel blocker LaCl₃ (Li *et al*., 2019a). Without LaCl₃ treatment, laser ablation induced a rapid primary Ca^2+^ wave within seconds, followed by secondary Ca^2+^ activities (**Fig. 5A**). In contrast, LaCl₃ treatment significantly abolished both primary and secondary Ca²⁺ responses following wounding, while a weak primary spike and secondary flashing activity were occasionally observed in wound-adjacent cells (**Fig. 5B, 5D, 5E, Video S9**). To further assess whether Ca^2+^ influx from the extracellular space is required, we applied the membrane-impermeable Ca^2+^ chelator, 1,2-bis(*o*-aminophenoxy) ethane-*N,N,N′,N′*-tetraacetic acid (BAPTA), to restrict apoplastic Ca^2+^ influx. Our results demonstrated that BAPTA application significantly suppressed the secondary Ca^2+^ flashing, while the primary signal was not effectively suppressed. (**Fig. 5C, 5D, 5E, Video S10**).

**Figure 5.**
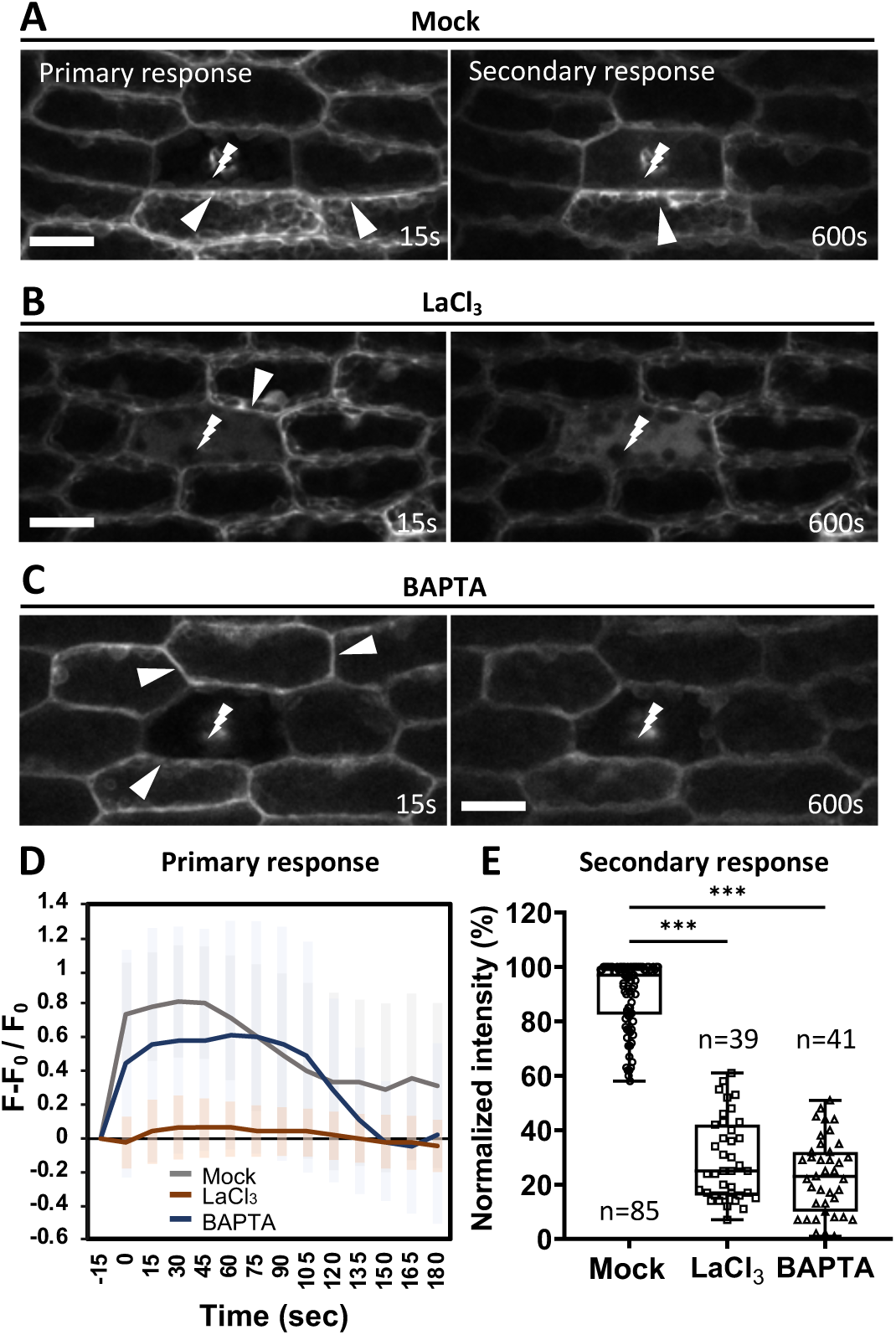
Regulated Ca^2+^ influx through plasma membrane channels activates wound-induced Ca^2+^ signaling. **(A)** Representative images showing laser-induced primary and secondary Ca^2+^ responses, indicated by arrowheads. The laser-ablated cell is marked by a lightning symbol. Scale bar = 20 µm. **(B)** and **(C)** Leaves excised from the GCaMP3 reporter line were pre-treated for 2 hours with either the Ca^2+^ channel blocker LaCl₃ (5 mM) **(B)** or the Ca^2+^ chelator BAPTA (0.5 mM) **(C)**, followed by laser ablation and time-lapse imaging. Under both treatments, only weak secondary Ca^2+^ activity was detected in neighboring cells. Scale bar = 20 µm. **(D)** Quantification of the primary Ca^2+^ spike in wound-adjacent cells. Mean fluorescence intensity was measured in wound-neighbor cells before and after wounding (t = 0 s). Wounding rapidly induced a transient Ca^2+^ spike, which was effectively suppressed by 5 mM LaCl₃ treatment. In contrast, suppression of the Ca^2+^ spike was less pronounced in BAPTA-treated samples. Number of cells =38 for mock, 19 for LaCl₃ treatment, 20 for BAPTA treatment. **(E)** Quantification of the secondary signals in wound-adjacent cells. For each treatment, fluorescence intensity was measured before and after treatment. The mean intensity before treatment was used as the normalization reference and set to 100%. Subsequent fluorescence values were expressed as percentages relative to the pre-treatment baseline. The number of quantified cells is indicated. *Student’s t-test, *** P<0.001*.

To assess the extent to which Ca^2+^ influx inhibitors suppress the wound response, we measured the intensity of the primary spike and the maximum intensity of the Ca^2+^ secondary responses in the absence or presence of these chemicals. For the primary spike, inhibition of Ca^2+^ channels and chelation of extracellular Ca^2+^ ions reduced the intensity to different degrees, and the signals were restricted to the wound-neighbor cells without spreading patterns (**Fig. 5D, Videos S9, S10**). For the secondary response, mean fluorescence intensity was measured before and after treatment. For normalization, the average intensity before treatment was defined as 100%, and fluorescence values after treatment were presented as percentages relative to the corresponding pre-treatment intensity. The result showed approximately 80% suppression of secondary Ca^2+^ signals when Ca^2+^ channels were blocked, or intercellular Ca^2+^ was chelated (**Fig. 5E**). Despite the significant suppression, weak primary and secondary responses can still be detected in wound-neighbor cells. We suspect that, in addition to apoplastic Ca^2+^ sources, Ca^2+^ stored in laser-target cells may be released and transported to wound-neighbor cells via plasmodesmata. Together, these results indicate that wound-induced Ca^2+^ signaling predominantly depends on Ca^2+^ influx from the extracellular space, suggesting Ca^2+^ influx occurs through regulated plasma membrane pathways rather than passive, non-selective diffusion resulting from membrane damage.

### Wound-induced Ca^2+^ signaling is required for rapid F-actin reorganization at wound sites

To investigate whether rapid Ca^2+^ influx contributes to other early wound responses, we examined the role of Ca^2+^ signaling in wound-induced F-actin reorganization. Our previous study has shown that F-actin undergoes a transient and localized accumulation at the wound-adjacent plasma membrane of neighboring cells following wounding (Huang *et al*., 2025). To test whether this F-actin response depends on rapid Ca^2+^ signaling, we applied the Ca^2+^ chelator BAPTA to suppress wound-induced Ca^2+^ influx. Under these conditions, F-actin accumulation at the wound-adjacent membrane was strongly reduced compared with untreated controls (**Fig. 6A, 6B**). Although weak F-actin enrichment was occasionally observed in a small number of neighboring cells, the magnitude and spatial restriction of this response were markedly diminished. In contrast, non-neighboring cells did not exhibit significant differences in F-actin organization between control and BAPTA-treated samples (**Fig. 6C**). In addition to the chelator, we also tested LaCl_3_ treatment and observed a similar inhibitory effect on F-actin accumulation in wound-neighbor cells. However, under the same laser ablation conditions, we noticed that LaCl_3_ treatment also suppressed membrane deformation (**Fig. S2**). Therefore, to exclude the possibility that the absence of an F-actin response results from impaired membrane deformation, we applied BAPTA treatment and verified membrane deformation in each experiment (N=4 independent experiments). These results indicate that Ca^2+^ influx is required for wound-induced F-actin reorganization specifically in wound-adjacent cells.

**Figure 6.**
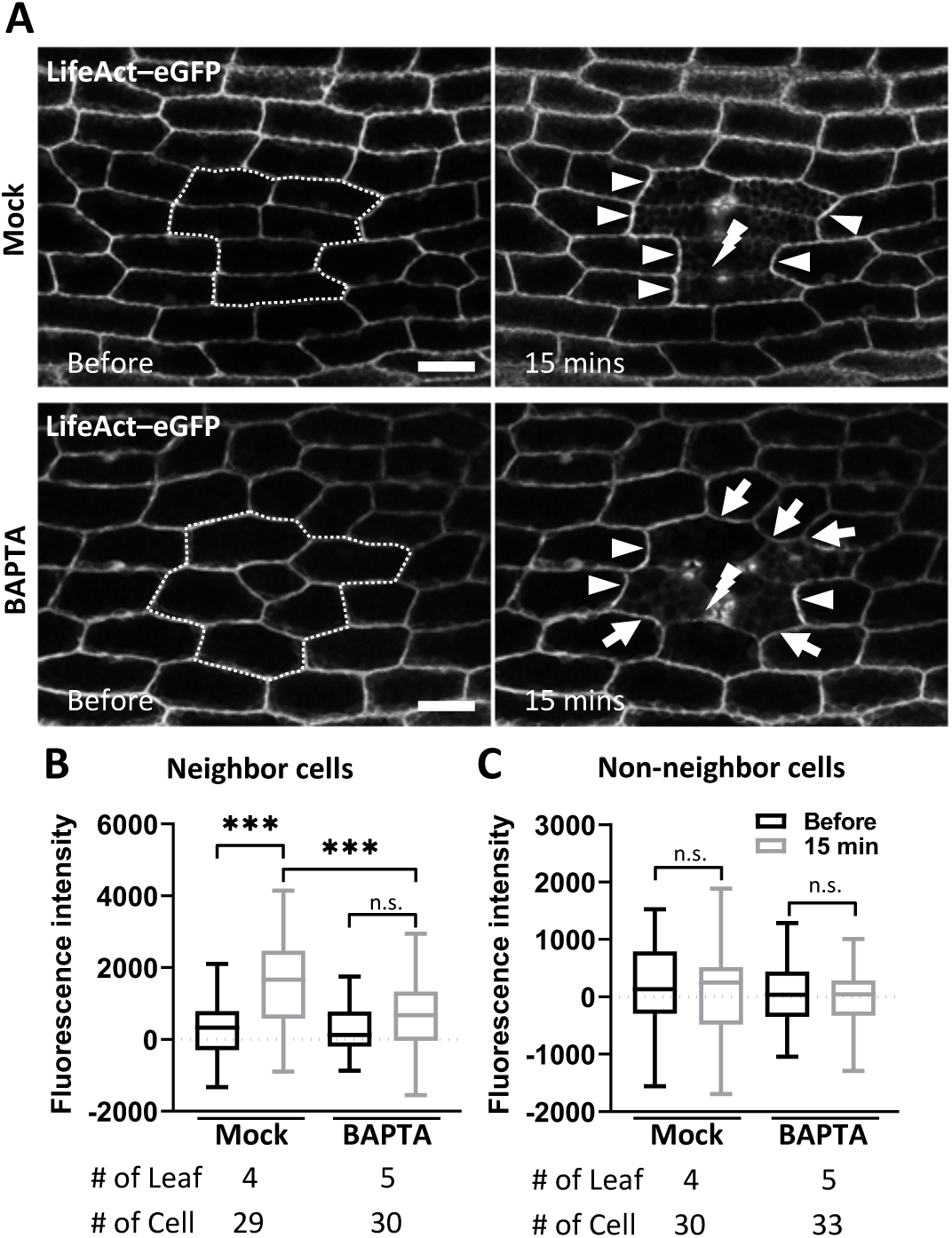
Wound-induced Ca^2+^ signaling is required for F-actin reorganization at wound-adjacent cells. **(A)** Representative images showing wound-induced F-actin reorganization visualized using the LifeAct–eGFP marker. Six-cell laser ablation was performed at *t* = 0 min. The ablated area is indicated by one lightning symbol and a dotted frame. In control samples, LifeAct–eGFP signals were enriched at the wound-adjacent plasma membrane of neighboring cells 15 minutes after ablation, with prominent accumulation in axially aligned cells (arrowheads). Lower panels show the same experimental setup following pre-treatment with the Ca^2+^ chelator BAPTA (0.5 mM, 2 hours). Under Ca^2+^ chelation, F-actin accumulation at the wound-adjacent membrane was strongly reduced. Arrowheads point to weak residual F-actin enrichment, and arrows mark wound-adjacent membrane lacking detectable F-actin accumulation. Scale bar = 20 µm. **(B)** and **(C)** Quantitative analysis of LifeAct-eGFP signals on the interface membrane of neighbor cells and non-neighbor cells at t=before and 15 minutes. *Student’s t-test, *** P<0.001.* n.s., not significant.

### Mechanosensitive Ca^2+^ signaling is essential for wound-induced cell regeneration

Next, we asked whether rapid wound-induced Ca^2+^ signaling is required for the initiation of cell reprogramming and regeneration. To address this, we blocked Ca^2+^ influx using a low concentration (0.1 mM) of LaCl₃ and examined both the initiation of cell reprogramming and subsequent regeneration efficiency. Here, we chose LaCl_3_ over BAPTA because, during long-term observation, BAPTA treatment caused cell condensation and cytotoxicity (**Fig. S3**). Although LaCl₃ treatment suppressed membrane deformation under standard laser ablation conditions, we found that a more severe wounding method, e.g., needle poking, as we performed in regeneration assays, was still able to cause membrane deformation in the presence of LaCl₃ (**Fig. S4**). Meanwhile, the Ca^2+^ flashing signal was sufficiently inhibited under this condition (**Video S11**).

Initiation of cell reprogramming during wound-induced regeneration has been characterized previously using the protonema-specific marker RM55. Activation of RM55 expression marks the reprogramming of gametophore cells into protonema stem cells (Ishikawa *et al*., 2011). We therefore used the same RM55 reporter line to assess whether inhibition of Ca^2+^ influx affects the onset of cell reprogramming following wounding. To increase the number of wounding sites on one excised leaf and observe the RM55 reporter, we performed needle poking as a wounding method in our regeneration assay. We first confirmed that Ca^2+^ flashing signals can be consistently observed under this wounding method (**Video S12**). In untreated samples, RM55 expression was robustly induced in cells surrounding the wound approximately 22 hours after wounding. In contrast, RM55 activation was undetectable in LaCl₃-treated samples (**Fig. 7A, 7B**). To note, the concentration of LaCl_3_ used in the long-term regeneration experiments (0.1 mM) was substantially lower than that used in the short-term assays (5 mM; **Fig. 5**). This lower concentration has been reported previously to cause only a mild delay in the growth of *P. patens* protonemal cells, indicating that cells remain viable and capable of slow but normal growth under these conditions (Bascom *et al*., 2018). To further assess potential effects on viability in our system, we performed Propidium Iodide staining on excised leaves. The results showed that, after 16 hours of LaCl₃ treatment, leaf cells remained alive (**Fig. S5**).

**Figure 7.**
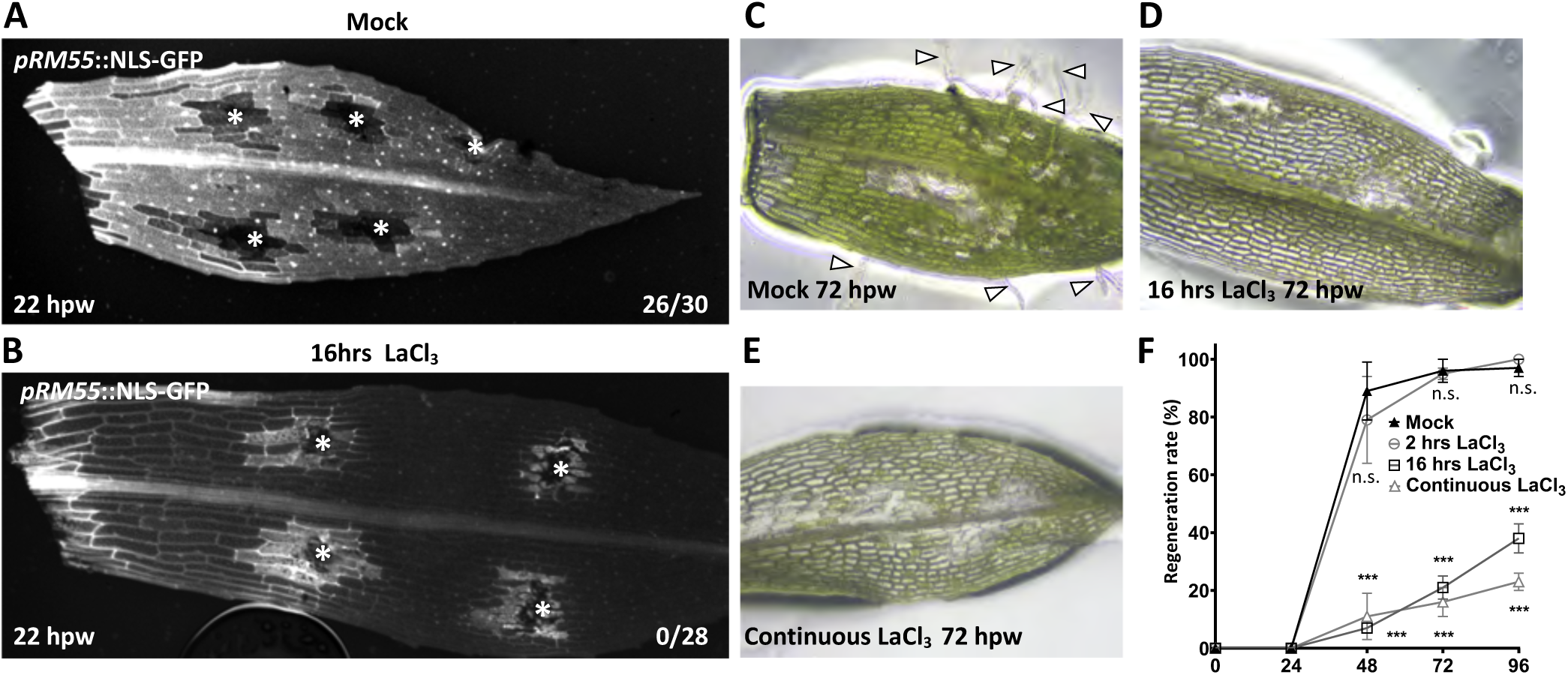
Mechanosensitive Ca^2+^ signaling is essential for wound-induced cell reprogramming and regeneration. **(A)** and **(B)** Representative images of the *pRM55*::NLS-GFP reporter line with or without LaCl_3_ treatment. *pRM55*::NLS-GFP is activated when gametophore cells reprogram into protonema stem cells, indicating the initiation of cell reprogramming. GFP is fused to a nuclear localization signal (NLS). **(A)** Reporter activation was detected at 22 hours post-wounding (hpw) in the absence of LaCl_3_ treatment. **(B)** Application of 0.1 mM LaCl₃ during the first 16 hours post-wounding. No detectable nuclear GFP signal was observed at 22 hpw. The blurry fluorescence surrounding wound sites was auto-fluorescence emitted by dead cells. Wound sites generated by needle injury are marked by asterisks. The number of wound sites exhibiting GFP-positive nuclei is indicated in the lower right corners and shown relative to the total number of wound sites. **(C–E)** Excised leaves of the LifeAct–eGFP line were cultured on a water plate (mock) **(C)**, an agar plate containing 0.1 mM LaCl₃ for 16 hours after wounding **(D)**, or an agar plate containing 0.1 mM LaCl₃ continuously **(E)**, and hollow arrowheads point to the regenerative outgrowing cells. **(F)** The regeneration event was monitored every 24 hours using light microscopy. Regeneration rate was quantified as the proportion of leaves exhibiting regeneration events, including cell outgrowth and regenerative division, relative to the total number of excised leaves. N=3-4 independent experiments, n≥30 leaves per experiment. *Student’s t-test, *** P<0.001,* n.s., not significant.

We next asked whether impaired initiation of cell reprogramming reduces the regenerative capacity. Given that secondary Ca^2+^ activity persists for up to 16 hours after wounding, we treated excised leaves with LaCl₃ continuously or for the first 16 hours following wounding **(Fig. 7C-7E**). In both conditions, regeneration was strongly suppressed. Most excised leaves grown on LaCl₃-containing plate failed to regenerate even 96 hours after wounding, in contrast to robust regeneration in control samples (**Fig. 7F**). Moreover, we tested whether the early Ca^2+^ spike is required for regeneration by transiently inhibiting Ca^2+^ signaling during the first two hours after wounding (**Fig. 7F**). Transient inhibition during this early window did not affect regeneration, suggesting that the primary Ca^2+^ spike may not be necessary to initiate cell fate reprogramming. Instead, these results elucidate that prolonged Ca^2+^ dynamics are specifically required to enable reprogramming and regeneration, while the early Ca^2+^ spike likely represents an immediate but functionally distinct wound response. Together, these results demonstrate that rapid wound-induced Ca^2+^ signaling, particularly prolonged secondary flashing activity, is essential for the initiation of cell reprogramming and successful regeneration.

## Materials and Methods

### Plant material and growth conditions

For the observation of cytoskeletal networks and reprogramming rates, *Physcomitrium patens*, Gransden strain, (Ashton and Cove, 1977) was used. Moss tissues were routinely grown on BCDAT (BCD medium contains 1 mM MgSO_4_, 10 mM KNO_3_, 45 µM FeSO_4_, 1.8mM KH_2_PO_4_ [pH 6.5 adjusted with KOH], and trace element solution (0.22 µM CuSO_4_, 0.19 µM ZnSO_4_, 10 µM H_3_BO_3_, 0.10 µM Na_2_MoO_4_, 2 µM MnCl_2_, 0.23 µM CoCl_2_, 0.17 µM KI); BCDAT is BCD medium with 1 mM CaCl_2_, and 5 mM diammonium (+)-tartrate) plates under 16-hour light/8-hour dark photoperiod at 25°C as described previously (Nishiyama *et al*., 2000).

For the Ca^2+^ signal observation, a GCaMP3 reporter line was made in a previous study. A circularly permuted green fluorescent protein(cpGFP) labeling was linked with M13 on Calmodulin driven by *Zea mays UBIQUITIN1* promoter, and a strategically employed *Porites porites* RFP as an expression control, driven by *Panicum virgatum POLYUBIQUITIN1* promoter (Kleist *et al*., 2017).

For F-actin observation, we utilized transgenic plants labeled F-actin and microtubules as previously described (Overdijk *et al*., 2016). In summary, a moss line that expresses a gene encoding mCherry fused to α-tubulin, driven by the rice actin promoter (McElroy *et al*., 1990) and integrated at the HB7 locus (Hiwatashi *et al*., 2008), was transformed with a LifeAct-eGFP construct (Vidali *et al*., 2009), thus making it a cytoskeleton double labeling line.

For the examination of cell reprogramming, we employed a cell reprogramming marker, RM55, which has an NLS-GFP signal driven by the protonema-specific promoter *pRM55*, and has been verified as a reprogramming marker in *P. patens* leaves (Ishikawa *et al*., 2011).

### Laser-induced moss leaf cell Calcium signal

Excised leaves from 4-week-old *P. p*atens were carefully transferred onto a slide containing ddH_2_O and covered with a coverslip. Precisely localized cell damage was induced using a high-resolution confocal laser scanning microscope (Zeiss LSM 900) equipped with 405, 488, and 561 nm lasers. Laser power was 100%, and the laser target area was shaped into circles. For the 6-cell ablation area, the target area was focused on the middle junction of the cells. The laser was conducted for 30 seconds (Zeiss LSM 900), and the irradiation was performed once. Bright-field or fluorescent imaging was used on all ablated cells to confirm whether a cell died. A cell is clearly dead when the shrinkage of the intracellular contents becomes visible. The other indicator was the emission of the strong auto-fluorescence from dead cells, which was detected with 561nm laser excitation.

### Propidium iodide staining

For PI (Propidium Iodide) staining, the excised leaves from 4-week-old *P. patens* were carefully transferred into a 400 *μl* of 10 *μ*g/ml PI in the dark for 15 minutes. After 15 minutes, the leaves were washed 2 times and transferred to a slide for imaging.

### Microscopy

For live-cell imaging, time-lapse images of the GCaMP3 were captured at 15- or 10-second intervals for 15 or 10 minutes, respectively. Images were acquired using Zeiss LSM900 with a 20x air objective. The GCaMP3 was excited with the 488 nm laser. Image processing, including image stacking and signal quantification, was performed in Fiji (Schindelin *et al*., 2012).

### Pharmacological Treatments

LaCl_3_ and BAPTA were used to inhibit the Ca^2+^ channel and chelate intercellular Ca^2+^ ions, respectively. The duration of each treatment was detailed in the main text. 5mM LaCl_3_ or 0.5 mM BAPTA was applied for the indicated duration as shown in **Fig. 5**. The chemicals were diluted in ddH_2_O, and leaves were submerged in the solution for 2 hours. For the long-term treatment, the excised leaves were placed on water plates with or without LaCl_3_ (0.1 mM) supplement for 16 hours and carefully transferred to another water plate for regeneration rate observation (Bascom *et al*., 2018).

### Hyper-osmotic and Hypo-osmotic test

For the hyper-osmotic treatment, a microfluidic chamber was used. Consists of one 24×50 mm and one 22×22 mm coverslip with a piece of 3M 467MP 2 mil Transfer Tape – 200MP Acrylic Adhesive sticker. The size of the sticker is 35 × 11 mm with a hollow section in the middle. The central cut-out is a continuous, symmetric constricted channel, which can be segmented into three functional regions along the long axis. The entrance region starts from 10 mm from the edge of the sticker, consists of a 4×4 mm square, and is followed by a trapezoid. The 4 mm of the trapezoid is gradually decreased to 0.7 mm within the distance of 14 mm, followed by the sample region. The sample region is a 2×2 mm square, followed by the exit region. The exit region consists of a rectangular gate with a 1×2.2 mm gate with the shorter side connect with sample region. At the other side of the rectangular gate is a 2.5×2.5 mm square for liquid removal. The mannitol concentration used in the hyper-osmotic test were 750 mM (Storti *et al*., 2018a). Hyper-osmotic wave and hypo-osmotic test were filmed at 10-second intervals.

### Measurement of reprogramming and regeneration rate

For the reprogramming initiation, wounded leaves collected from the RM55 line were placed on a BCDAT medium and observed daily for one week using a Zeiss upright fluorescence microscope with a GFP filter. Images were acquired with a 10x air objective and 1024 x 1024-pixel resolution. The regeneration rate was measured as previously described (Huang *et al*., 2025). In brief, cell regeneration events were observed and recorded as individual occurrences. The cell regeneration rate at each time point was calculated by dividing the number of leaves with at least one regeneration event, including cell outgrowth and regenerative division, by the total number of leaves.

### Image data analysis

GCaMP3 fluorescence intensity was quantified using Fiji (Schindelin *et al*., 2012). For the flashing and aggregation patterns of the GCaMP3 signal on the plasma membrane, a line with a 5-pixel thickness was plotted along the plasma membrane indicated with arrowheads, and the mean intensity along the line overtime was shown in kymograph (**Fig. 1**). Statistical analysis was performed using *Student’s t-test* (**Fig. 5, 6, 7**), and results were presented as mean ± standard deviation. Statistical significance was set at *p < 0.05*.

## Discussion

### Spatiotemporal framework of wound-induced Ca^2+^ signaling in *P. patens*

In this study, we identified a rapid, wound-induced Ca^2+^ wave that spread from the wound-neighboring cells and decayed within minutes (**Fig. 1**), resembling mechanically induced Ca^2+^ responses reported in other plant systems and in *P. patens* under touch stimulation (Kleist *et al*., 2017; Zhou *et al*., 2025). Strikingly, this primary response was followed by a previously uncharacterized secondary Ca^2+^ activity specifically in wound-adjacent cells, exhibiting as either stochastic flashing or polarized aggregation. Although the primary wave was consistently observed in single-cell and 6-cell ablation sites with no discernible differences, the aggregated form of the secondary activity occurred predominantly in single-cell ablation. We speculate that wound severity must exceed a certain threshold to induce the flashing response, whereas single-cell ablation may not always generate sufficient damage to trigger flashing and instead results in a localized aggregation signal. This aggregation may arise from the accumulation of Ca^2+^ released from the ablated cell. In contrast, the flashing response may reflect a combined process involving both extracellular Ca^2+^ influx and the propagation or redistribution of Ca^2+^ originating from damaged cells.

Unlike canonical Ca^2+^ spikes, these secondary responses were non-periodic, spatially heterogeneous, and persisted for hours (**Fig. 2**). To our knowledge, Ca^2+^ signaling events lasting beyond 10-15 minutes have never been reported in response to mechanical wounding in plants (Tian *et al*., 2020; Toyota *et al*., 2018). Thus, our data reveal a unique prolonged Ca^2+^ signaling signature associated with wound perception in *P. patens*.

### The role of localized membrane deformation in shaping Ca^2^**^+^** dynamics

A central question in plant mechanobiology is how transient physical stimuli are translated into sustained developmental signals. Our results provide a new perspective by distinguishing between initial perception and long-term maintenance of Ca^2+^ signatures. We observed that both localized wounding by poking or laser and generalized osmotic stress through plasmolysis induce an immediate Ca^2+^ spike, suggesting that various forms of membrane-wall disturbance can activate mechanosensitive ion channels. However, a substantial divergence occurs in the spatial dimension; only the wound-induced membrane deformation surrounding the wound site is associated with a local flashing signature. This suggests that the geometry and intensity of membrane deformation may encode specific information. Unlike the uniform membrane shrinkage during plasmolysis, which failed to generate spatial specificity for the Ca^2+^ flashing, the localized poking creates an asymmetric mechanical gradient. We propose that this specific physical state might be necessary to keep mechanosensitive complexes (such as MSLs or MCAs) in a “sensitized” state or to trigger a positive feedback loop involving ROS or cell-wall-derived DAMPs (Damage-Associated Molecular Patterns) (Hamilton *et al*., 2015; Marcec *et al*., 2019).

While we cannot rule out the involvement of downstream chemical ligands, the precise spatial co-localization of the prolonged signature with the deformed membrane area strongly supports the idea that localized physical strain is a primary requirement for the reprogramming-associated Ca^2+^ code. By maintaining this “calcium window” for multiple hours, the window between wounding and the activation of master regulators (Ishikawa *et al*., 2019), the cell may achieve the necessary threshold for the epigenetic and transcriptional resetting required for cell reprogramming.

### Extracellular Ca^2+^ is the primary source of wound-induced Ca^2+^ signals

Pharmacological inhibition supports the idea that regulated Ca^2+^ influx across the plasma membrane is the major source of wound-induced Ca^2+^ signaling. The inhibition of the primary Ca^2+^ wave by these treatments indicates that extracellular Ca^2+^ in the apoplast is the dominant source of the initial response. Weak secondary activity was occasionally detected, suggesting that intracellular Ca^2+^ stores may contribute to sustaining the flash signaling. While PIEZO functions as a plasma membrane-localized mechanosensitive Ca^2+^ channel in animals and *Arabidopsis*, *P. patens* PIEZO localizes to the vacuolar membrane and mediates Ca^2+^ release into the cytoplasm (Radin *et al*., 2021; Xiao, 2024). Whether vacuolar Ca^2+^ release contributes to prolonged secondary Ca^2+^ activity will require genetic analysis of *piezo* mutants and direct observation of vacuolar dynamics following wounding.

### Mechanosensitive Ca^2+^ signaling is required for wound-induced F-actin reorganization

Ca^2+^ signaling is known to regulate actin dynamics, often displaying a negative correlation with F-actin accumulation in tip-growing cells (Bascom *et al*., 2018; Cárdenas *et al*., 2008; Ryken *et al*., 2025). Ca^2+^-dependent actin-binding proteins directly translate Ca^2+^ fluctuations into cytoskeletal rearrangements (Yang *et al*., 2014; Yokota *et al*., 2000). In contrast to tip growth, our results revealed that inhibition of Ca^2+^ influx significantly reduced wound-induced F-actin accumulation at the wound-adjacent membrane, particularly in axial neighboring cells. Notably, 6-cell ablation predominantly elicited flashing Ca^2+^ responses, which consistently coincided with robust F-actin accumulation. These observations suggest that Ca^2+^ flashing, rather than aggregation, is required for F-actin reorganization during wound response. Although flashing Ca^2+^ signals exhibited distinct patterns from F-actin accumulation, they remained confined to wound-neighboring cells, where Ca^2+^ may play a role in coordinating localized cytoskeletal remodeling.

### Mechanosensitive Ca^2+^ signaling is associated with wound-induced regeneration

Rapid wound signaling has been extensively studied in the context of biotic and abiotic stress responses, such as pattern-triggered immunity and herbivore attack, where Ca^2+^ signaling activates jasmonate pathways and cell wall reinforcement to counteract damage (Bacete and Hamann, 2020; Zhao and Long, 2022). Our findings extend the functional scope of Ca^2+^ signaling to tissue regeneration.

Blocking Ca^2+^ influx abolished activation of the reprogramming marker and severely suppressed regeneration, suggesting that prolonged Ca^2+^ signaling is closely associated with the initiation of cell reprogramming. We acknowledge that prolonged LaCl₃ treatment may also produce pleiotropic effects that indirectly affect regeneration. However, continuous treatment with 0.1 mM LaCl₃ for seven days has previously been reported to cause only a mild delay in protonemal growth in *P. patens* (Bascom *et al*., 2018), and our viability assays showed that 16-hour LaCl₃ treatment did not result in detectable cell death (**Fig. S5**). These observations suggest that inhibition of Ca^2+^ influx, either directly or indirectly, interferes with the reprogramming process following wounding. How Ca^2+^ signaling interacts with other regeneration-associated pathways, including jasmonate signaling, cell wall remodeling, and chromatin modification, remains to be determined.

Mechanical forces have been proposed as upstream regulators of cell fate specification in diverse developmental contexts, including the shoot apical meristem and ovule primordia (Hernandez-Lagana *et al*., 2021; Landrein *et al*., 2015). In line with this, our study provides direct evidence that mechanical force-triggered Ca^2+^ signaling is a critical mediator connecting tissue damage to cell reprogramming and regeneration. Elucidating how this pathway is conserved or diversified across plant lineages and tissue types will be an important direction for future research.

### A working model for wound-induced Ca^2+^ signaling and regeneration

Based on our findings, we speculate that to trigger wound-induced cell reprogramming, the wound severity must exceed a certain threshold to induce the secondary flashing response, which may involve both extracellular Ca^2+^ influx and the release of Ca^2+^ from damaged or dead cells. Notably, the flashing signal persists for several hours within the temporal window between wounding and the onset of cell reprogramming, raising the possibility that these prolonged Ca^2+^ dynamics may contribute to transcriptional or epigenetic reprogramming during regeneration. In addition, flashing Ca^2+^ activity was associated with enhanced F-actin reorganization, suggesting a role in coordinating early wound responses at the cellular level. Together, these observations support a model in which flashing Ca^2+^ signaling acts as a mechanochemical cue linking physical injury to downstream cellular remodeling and regenerative responses.

## Supplemental data

The following supplementary data are available at JXB online.

Fig. S1. Laser ablation under hyper-osmotic stress fails to induce plasma membrane deformation.

Fig. S2. High concentration of LaCl_3_ inhibits laser-induced PM deformation.

Fig. S3. BAPTA treatment causes cell damage.

Fig. S4. LaCl_3_ treatment does not suppress the plasma membrane deformation caused by needle poking.

Fig. S5. Cell viability test with long-term LaCl_3_ treatment.

Fig. S6. Quantification of Ca^2+^ signals before and after recovering from LaCl_3_ treatment.

Video S1. Ca^2+^ primary wave in response to wounding.

Video S2. Secondary flashing signal in 1-cell ablation.

Video S3. Secondary aggregation signal in 1-cell ablation.

Video S4. Secondary flashing signal in 6-cell ablation.

Video S5. Ca^2+^ flashing signal at leaf cutting edge.

Video S6. No flashing signals on unwound leaf.

Video S7. Ca^2+^ spike under hypo-osmotic stress.

Video S8. 8 hours recovery from plasmolysis.

Video S9. Inhibition of Ca^2+^ signals with LaCl_3_ treatment.

Video S10. Inhibition of Ca^2+^ signals with BAPTA treatment.

Video S11. LaCl_3_ inhibits needle poking-induced Ca^2+^ signals.

Video S12. Needle poking induces Ca^2+^ flashing signals.

## Acknowledgments

The authors sincerely thank Dr. Kazimierz Trebacz’s kind sharing of the critical material GCaMP3 line, Dr. Ya-Yu Chiang for providing the microfluidic device, Dr. Mitsuyasu Hasebe for sharing the RM55 reporter line, and Dr. Barbara Kloeckener Gruissem’s time and efforts in critical reading and providing constructive advice on the manuscript.

## Author contributions

H.T. and H.A.W. initiated the project. H.A.W. carried out the experiments and performed quantitative analysis with Y.C.Y. support. H.T. wrote the manuscript.

## Data Availability

The data underlying this article are available in the article and its online supplementary material.

## Reference

Ashton N, Cove D. 1977. The isolation and preliminary characterisation of auxotrophic and analogue resistant mutants of the moss, Physcomitrella patens. Molecular and General Genetics MGG 154, 87–95.

Bacete L, Hamann T. 2020. The role of mechanoperception in plant cell wall integrity maintenance. Plants 9, 574.

Bascom CS, Winship LJ, Bezanilla M. 2018. Simultaneous imaging and functional studies reveal a tight correlation between calcium and actin networks. Proceedings of the National Academy of Sciences 115, E2869–E2878.

Bascom Jr CS, Winship LJ, Bezanilla M. 2018. Simultaneous imaging and functional studies reveal a tight correlation between calcium and actin networks. Proceedings of the National Academy of Sciences 115, E2869–E2878.

Birnbaum KD, Alvarado AS. 2008. Slicing across kingdoms: regeneration in plants and animals. Cell 132, 697–710.

Cárdenas L, Lovy-Wheeler A, Kunkel JG, Hepler PK. 2008. Pollen tube growth oscillations and intracellular calcium levels are reversibly modulated by actin polymerization. Plant Physiology 146, 1611–1621.

Fan X, Hou J, Chen X, Chaudhry F, Staiger CJ, Ren H. 2004. Identification and characterization of a Ca2+-dependent actin filament-severing protein from lily pollen. Plant Physiology 136, 3979–3989.

Guichard M, Thomine S, Frachisse J-M. 2022. Mechanotransduction in the spotlight of mechano-sensitive channels. Current opinion in plant biology 68, 102252.

Hamant O, Heisler MG, Jonsson H, Krupinski P, Uyttewaal M, Bokov P, Corson F, Sahlin P, Boudaoud A, Meyerowitz EM. 2008. Developmental patterning by mechanical signals in Arabidopsis. science 322, 1650–1655.

Hamilton ES, Schlegel AM, Haswell ES. 2015. United in Diversity: Mechanosensitive Ion Channels in Plants. Annual Review of Plant Biology 66, 113–137.

He L, Tao J, Maity D, Si F, Wu Y, Wu T, Prasath V, Wirtz D, Sun SX. 2018. Role of membrane-tension gated Ca2+ flux in cell mechanosensation. Journal of cell science 131, jcs208470.

Hernandez-Lagana E, Mosca G, Mendocilla-Sato E, Pires N, Frey A, Giraldo-Fonseca A, Michaud C, Grossniklaus U, Hamant O, Godin C. 2021. Organ geometry channels reproductive cell fate in the Arabidopsis ovule primordium. elife 10, e66031.

Hiwatashi Y, Obara M, Sato Y, Fujita T, Murata T, Hasebe M. 2008. Kinesins are indispensable for interdigitation of phragmoplast microtubules in the moss Physcomitrella patens. The Plant Cell 20, 3094–3106.

Hoermayer L, Montesinos JC, Trozzi N, Spona L, Yoshida S, Marhava P, Caballero-Mancebo S, Benkova E, Heisenberg C-P, Dagdas Y. 2024. Mechanical forces in plant tissue matrix orient cell divisions via microtubule stabilization. Developmental Cell 59, 1333–1344. e1334.

Huang YT, Yen YC, Vermeer JE, Willemsen V, Tang H. 2025. F-Actin Polarization and Microtubule Integrity Direct Regeneration Patterns and Polarity of Cell Outgrowth in Wound-Induced Reprogramming. Plant, Cell & Environment.

Ikeuchi M, Favero DS, Sakamoto Y, Iwase A, Coleman D, Rymen B, Sugimoto K. 2019. Molecular Mechanisms of Plant Regeneration. Annual Review of Plant Biology 70, 377–406.

Ishikawa M, Morishita M, Higuchi Y, Ichikawa S, Ishikawa T, Nishiyama T, Kabeya Y, Hiwatashi Y, Kurata T, Kubo M. 2019. Physcomitrella STEMIN transcription factor induces stem cell formation with epigenetic reprogramming. Nature Plants 5, 681–690.

Ishikawa M, Murata T, Sato Y, Nishiyama T, Hiwatashi Y, Imai A, Kimura M, Sugimoto N, Akita A, Oguri Y. 2011. Physcomitrella cyclin-dependent kinase A links cell cycle reactivation to other cellular changes during reprogramming of leaf cells. The Plant Cell 23, 2924–2938.

Kleist TJ, Cartwright HN, Perera AM, Christianson ML, Lemaux PG, Luan S. 2017. Genetically encoded calcium indicators for fluorescence imaging in the moss Physcomitrella: GCaMP3 provides a bright new look. Plant Biotechnol J 15, 1235–1237.

Koselski M, Hoernstein SNW, Wasko P, Reski R, Trebacz K. 2023. Long-distance electrical and calcium signals evoked by hydrogen peroxide in physcomitrella. Plant and Cell Physiology 64, 880–892.

Kurusu T, Kuchitsu K, Nakano M, Nakayama Y, Iida H. 2013. Plant mechanosensing and Ca2+ transport. Trends in plant science 18, 227–233.

Landrein B, Kiss A, Sassi M, Chauvet A, Das P, Cortizo M, Laufs P, Takeda S, Aida M, Traas J. 2015. Mechanical stress contributes to the expression of the STM homeobox gene in Arabidopsis shoot meristems. elife 4, e07811.

Li T, Yan A, Bhatia N, Altinok A, Afik E, Durand-Smet P, Tarr PT, Schroeder JI, Heisler MG, Meyerowitz EM. 2019a. Calcium signals are necessary to establish auxin transporter polarity in a plant stem cell niche. Nature Communications 10, 726.

Li T, Yan A, Bhatia N, Altinok A, Afik E, Durand-Smet P, Tarr PT, Schroeder JI, Heisler MG, Meyerowitz EM. 2019b. Calcium signals are necessary to establish auxin transporter polarity in a plant stem cell niche. Nat Commun 10, 726.

Marcec MJ, Gilroy S, Poovaiah B, Tanaka K. 2019. Mutual interplay of Ca2+ and ROS signaling in plant immune response. Plant Science 283, 343–354.

McElroy D, Zhang W, Cao J, Wu R. 1990. Isolation of an efficient actin promoter for use in rice transformation. The Plant Cell 2, 163–171.

Nishiyama T, Hiwatashi Y, Sakakibara K, Kato M, Hasebe M. 2000. Tagged mutagenesis and gene-trap in the moss, Physcomitrella patens by shuttle mutagenesis. DNA research 7, 9–17.

Overdijk EJR, De Keijzer J, De Groot D, Schoina C, Bouwmeester K, Ketelaar T, Govers F. 2016. Interaction between the moss Physcomitrella patens and Phytophthora: a novel pathosystem for live-cell imaging of subcellular defence. Journal of Microscopy 263, 171–180.

Radin I, Richardson RA, Coomey JH, Weiner ER, Bascom CS, Li T, Bezanilla M, Haswell ES. 2021. Plant PIEZO homologs modulate vacuole morphology during tip growth. science 373, 586–590.

Ryken SE, Wu S-Z, Lee ML, Greig MM, Recto NM, Chang Stauffer S, Bascom CS, Jr., Kramer EM, Bezanilla M. 2025. Autoinhibitory calcium ATPases regulate the calcium gradient required for rapid polarized growth. Journal of Cell Biology 225.

Sakakibara K, Reisewitz P, Aoyama T, Friedrich T, Ando S, Sato Y, Tamada Y, Nishiyama T, Hiwatashi Y, Kurata T. 2014. WOX13-like genes are required for reprogramming of leaf and protoplast cells into stem cells in the moss Physcomitrella patens. Development 141, 1660–1670.

Schindelin J, Arganda-Carreras I, Frise E, Kaynig V, Longair M, Pietzsch T, Preibisch S, Rueden C, Saalfeld S, Schmid B. 2012. Fiji: an open-source platform for biological-image analysis. Nature methods 9, 676–682.

Storti M, Costa A, Golin S, Zottini M, Morosinotto T, Alboresi A. 2018a. Systemic Calcium Wave Propagation in Physcomitrella patens. Plant and Cell Physiology.

Storti M, Costa A, Golin S, Zottini M, Morosinotto T, Alboresi A. 2018b. Systemic calcium wave propagation in Physcomitrella patens. Plant and Cell Physiology 59, 1377–1384.

Suda H, Toyota M. 2022. Integration of long-range signals in plants: A model for wound-induced Ca2+, electrical, ROS, and glutamate waves. Current opinion in plant biology 69, 102270.

Sugimoto K, Temman H, Kadokura S, Matsunaga S. 2019. To regenerate or not to regenerate: factors that drive plant regeneration. Current opinion in plant biology 47, 138–150.

Tian W, Wang C, Gao Q, Li L, Luan S. 2020. Calcium spikes, waves and oscillations in plant development and biotic interactions. Nature Plants 6, 750–759.

Toyota M, Spencer D, Sawai-Toyota S, Jiaqi W, Zhang T, Koo AJ, Howe GA, Gilroy S. 2018. Glutamate triggers long-distance, calcium-based plant defense signaling. science 361, 1112–1115.

Vidali L, Van Gisbergen PA, Guérin C, Franco P, Li M, Burkart GM, Augustine RC, Blanchoin L, Bezanilla M. 2009. Rapid formin-mediated actin-filament elongation is essential for polarized plant cell growth. Proceedings of the National Academy of Sciences 106, 13341–13346.

Watanabe K, Hashimoto K, Hasegawa K, Shindo H, Tsuruda Y, Kupisz K, Koselski M, Wasko P, Trebacz K, Kuchitsu K. 2024. Rapid propagation of Ca2+ waves and electrical signals in the liverwort Marchantia polymorpha. Plant and Cell Physiology 65, 660–670.

Xiao B. 2024. Mechanisms of mechanotransduction and physiological roles of PIEZO channels. Nature Reviews Molecular Cell Biology 25, 886–903.

Yang X, Wang S-S, Wang M, Qiao Z, Bao C-C, Zhang W. 2014. Arabidopsis thaliana calmodulin-like protein CML24 regulates pollen tube growth by modulating the actin cytoskeleton and controlling the cytosolic Ca2+ concentration. Plant molecular biology 86, 225–236.

Yokota E, Muto S, Shimmen T. 2000. Calcium-calmodulin suppresses the filamentous actin-binding activity of a 135-kilodalton actin-bundling protein isolated from lily pollen tubes. Plant Physiology 123, 645–654.

Zhao F, Long Y. 2022. Mechanosensing, from forces to structures. Front Plant Sci 13, 1060018.

Zhou X, Li B, Li J, Sun Y, Xie R, Higashiyama T, Xiao S, Xin G, Su S. 2025. A mechanosensitive ion channel controls touch-triggered stigma movement through manipulation of calcium signature in Torenia. Nature Communications 16, 6296.

