## Supplemental figures for "Mechanosensing activates flashing Ca^2+^ dynamics associated with cell regeneration in *Physcomitrium patens*"

**Supplemental Figure 1. Laser ablation under hyper-osmotic stress fails to induce plasma membrane deformation.**

t = Before t = 600s

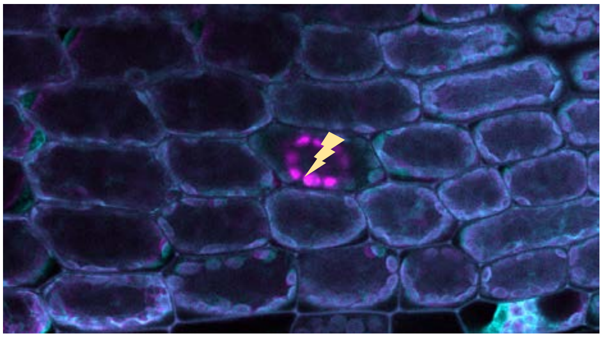

t = Before t = 900s

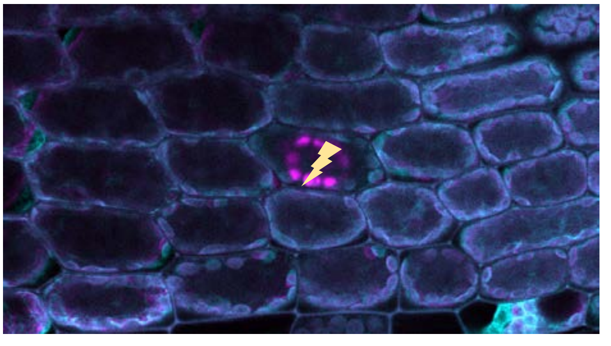

**Supplemental Figure 1. Laser ablation under hyper-osmotic stress fails to induce plasma membrane deformation.**
Excised leaves from GCaMP3 reporter lines were treated with mannitol for more than 20 min to induce hyperosmotic stress before laser ablation. Under these conditions, plasmolyzed cells showed no detectable change in cell shrinkage following ablation. Images acquired before and after laser treatment were pseudocolored in cyan and magenta, respectively. The merged images showed nearly complete overlap, indicating that laser ablation did not induce detectable plasma membrane deformation under hyperosmotic conditions.

**Supplemental Figure 2. High concentration of $LaCl_3$ inhibits laser-induced PM deformation.**

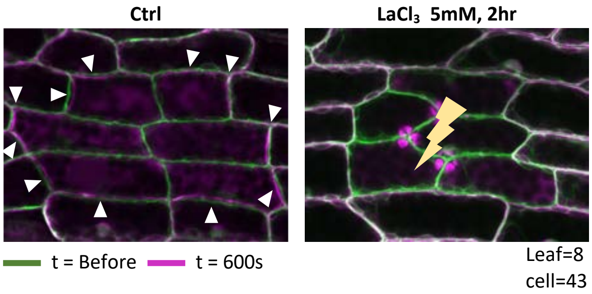


**Supplemental Figure 2. High concentration of $LaCl_3$ inhibits laser-induced PM deformation.**
Under growth medium, laser ablation of 6 cells induces clear plasma membrane (PM) deformation as indicated by arrowheads, while the 5 mM $LaCl_3$ supplement inhibits laser-induced PM deformation.

**Supplemental Figure 3. BAPTA treatment causes cell damage.**

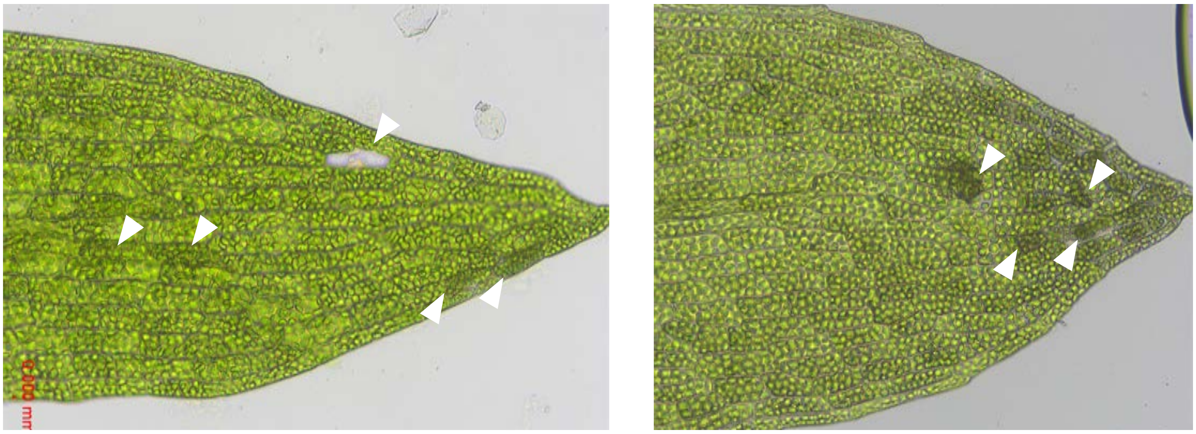

**Supplemental Figure 3. BAPTA treatment causes severe cell damage.**
Excised leaves were treated in growth medium supplied with 1mM BAPTA for 4 hours, which caused cell condensation and random death as indicated by white arrowheads.

**Supplemental Figure 4. $LaCl_3$ treatment does not suppress the plasma membrane deformation caused by needle poking.**

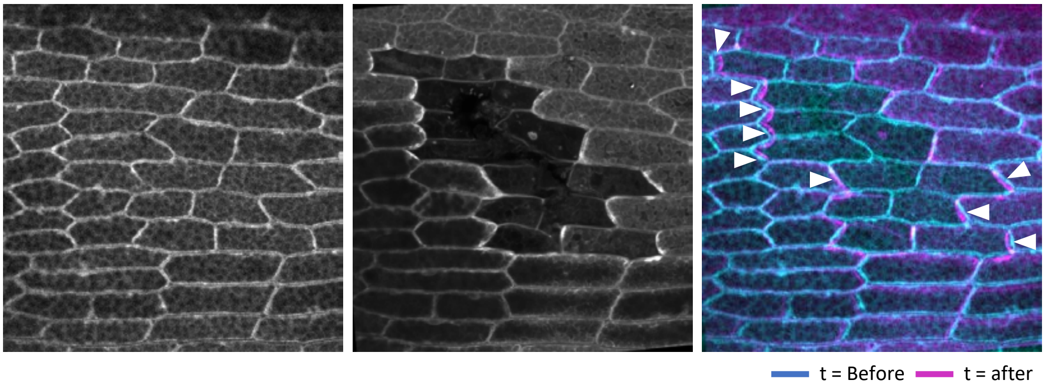


**Supplemental Figure 4. $LaCl_3$ treatment does not suppress the plasma membrane deformation caused by needle poking.**
Excised leaves from LifeaAct-eGFP marker line were imaged before treatment (left panel), followed by needle poking and 0.1 mM $LaCl_3$ treatment for 2 hours. The leaves were imaged again 2 hours after poking (middle panel). The images taken before and after poking were pseudocolored in cyan and magenta. The plasma membrane deformation caused by wounding was presented with overlaid images and indicated with white arrowheads. N=20 poking area.

**Supplemental Figure 5. Cell viability test with long-term $LaCl_3$ treatment.**

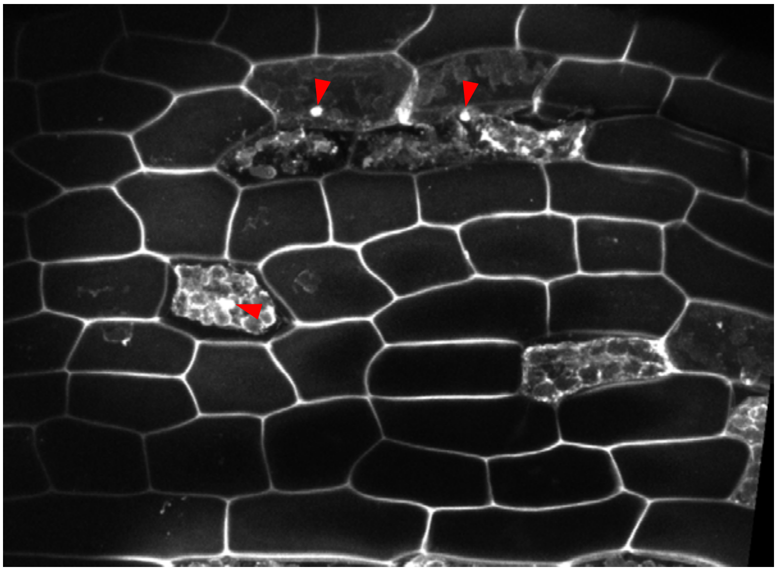

**Supplemental Figure 5. Cell viability test with long-term $LaCl_3$ treatment.**
The excised leaf was treated with 0.1 mM $LaCl_3$ for 16 hours, and stained with Propidium Iodide for 20 minutes. Only a few dead cells were stained with a bright nuclear signal, indicated by red arrowheads, demonstrating cell death. Most cells were stained only on the cell wall, indicating their viability.

**Supplemental Figure 6. Quantification of $Ca^{2+}$ signals before and after recovering from $LaCl_3$ treatment.**

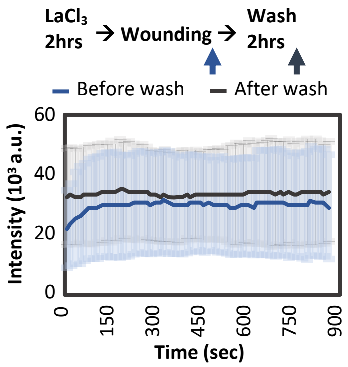


**Supplemental Figure 6. Quantification of $Ca^{2+}$ signals before and after recovering from $LaCl_3$ treatment.** Mean intensity across the membrane of wound-neighbor cells was measured over time. The 900s time-lapse imaging for before-wash and after-wash samples was acquired at the indicated time points by blue and gray arrows. n=25 cells for the before-wash sample; n=24 for the after-wash sample.
